## Supplemental Data for "CCDC113 stabilizes sperm axoneme and head-tail coupling apparatus to ensure male fertility"

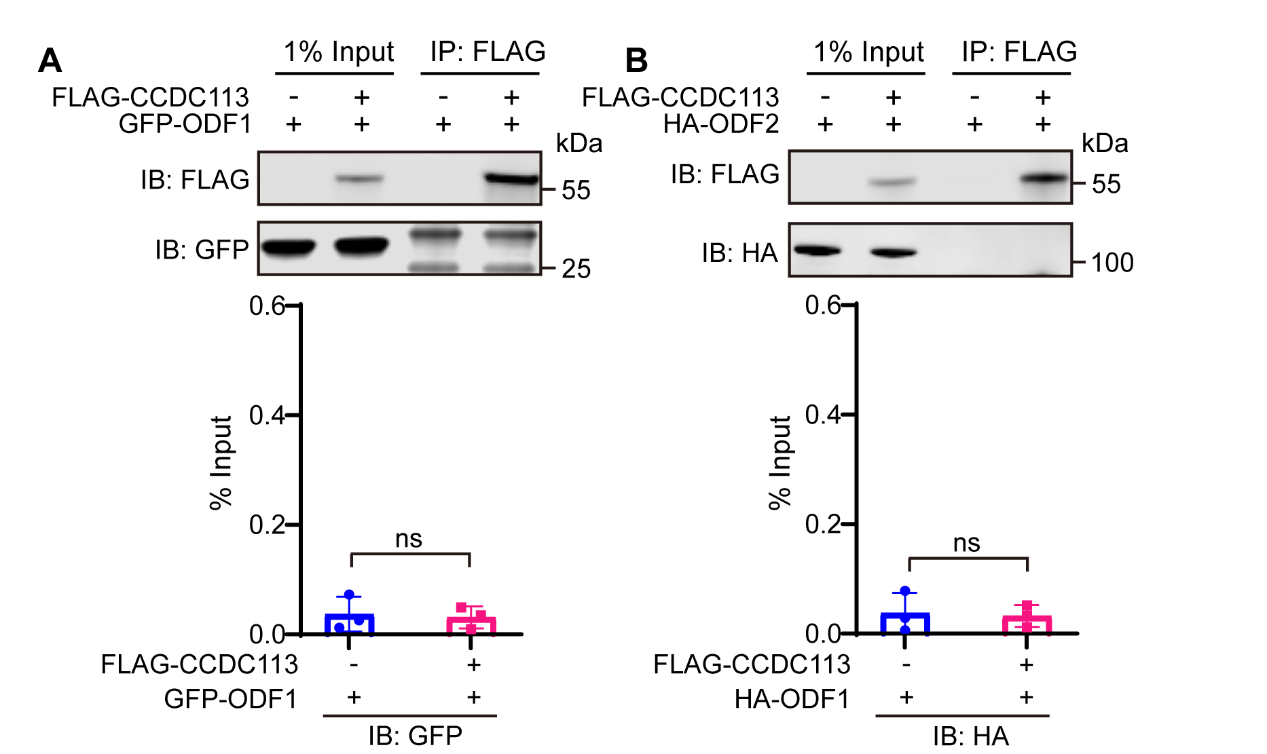

**Figure 1-figure supplement 1. CCDC113 could not bind to ODF1 and ODF2.**

(A-B) CCDC113 was expressed alone or co-expressed with ODF1 (A) or ODF2 (B) in HEK293T cells, and the interactions between CCDC113 and ODF1 or ODF2 were examined by co-immunoprecipitation. IB: immunoblotting; IP: immunoprecipitation. The % Input is displayed below the corresponding figures for quantification. *n* = 3 independent experiments. Data are presented as mean ± SD; ns indicates no significant difference.

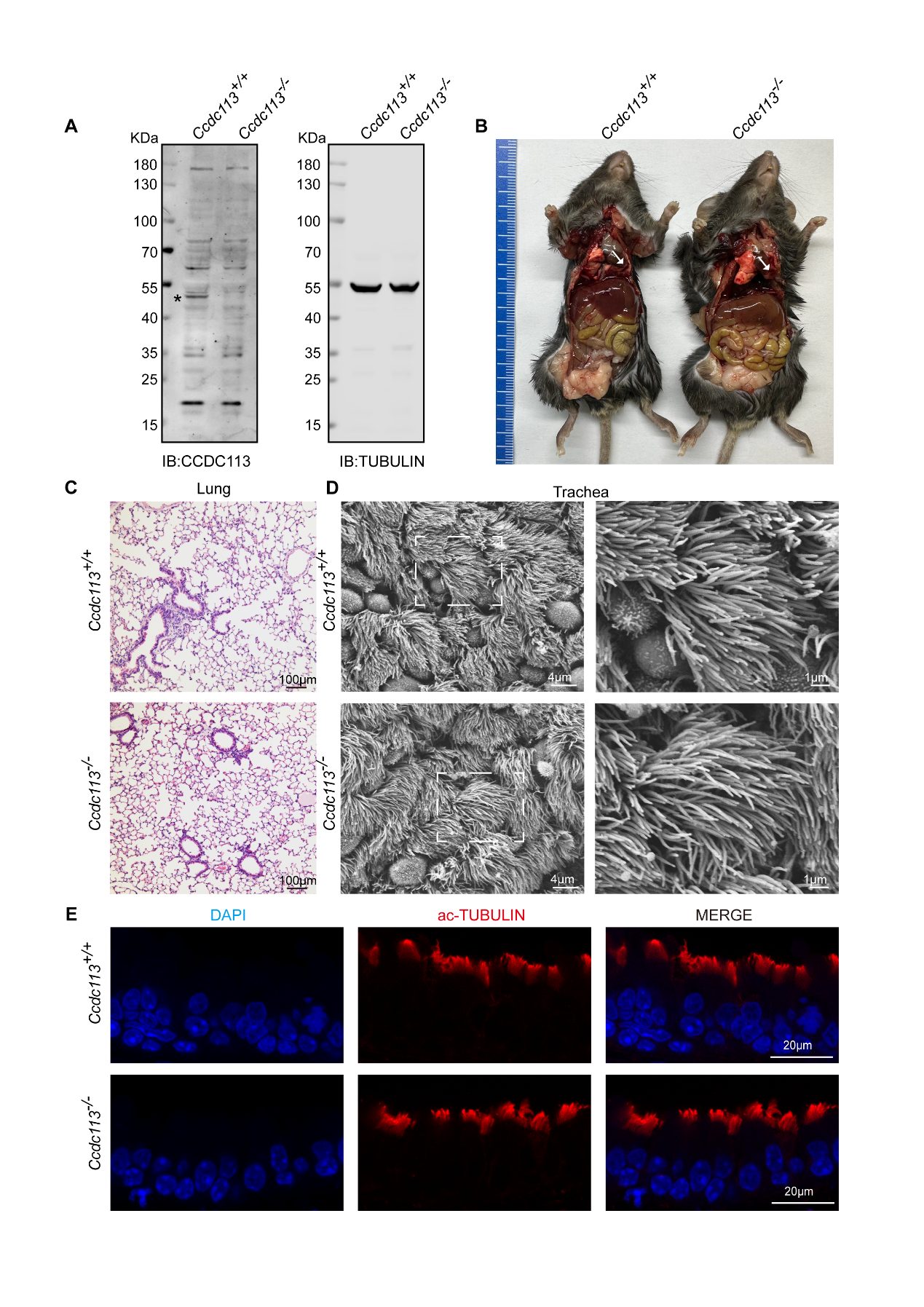

**Figure 2-figure supplement 1.** ***Ccdc113^–/–^* mice did not exhibit the ciliopathies, such as hydrocephalus, situs inversus, and abnormal ciliogenesis of tracheal cilia*.*** (A) Immunoblotting of CCDC113 in *Ccdc113^+/+^* and *Ccdc113^–/–^* testes, with TUBULIN serving as the loading control. An asterisk indicates the CCDC113 band. IB: immunoblotting. (B) The *Ccdc113^–/–^* mice did not exhibit hydrocephalus or left-right asymmetry defects. (C) Histology staining of the lung from *Ccdc113^+/+^* and *Ccdc113^-/-^* mice. (D) Scanning electron micrography of *Ccdc113^+/+^* and *Ccdc113^-/-^* tracheal epithelium at low and high magnifications of the boxed areas. (E) Immunofluorescence analysis of acetylated-tubulin (red) from *Ccdc113^+/+^* and *Ccdc113^-/-^* trachea cilia.

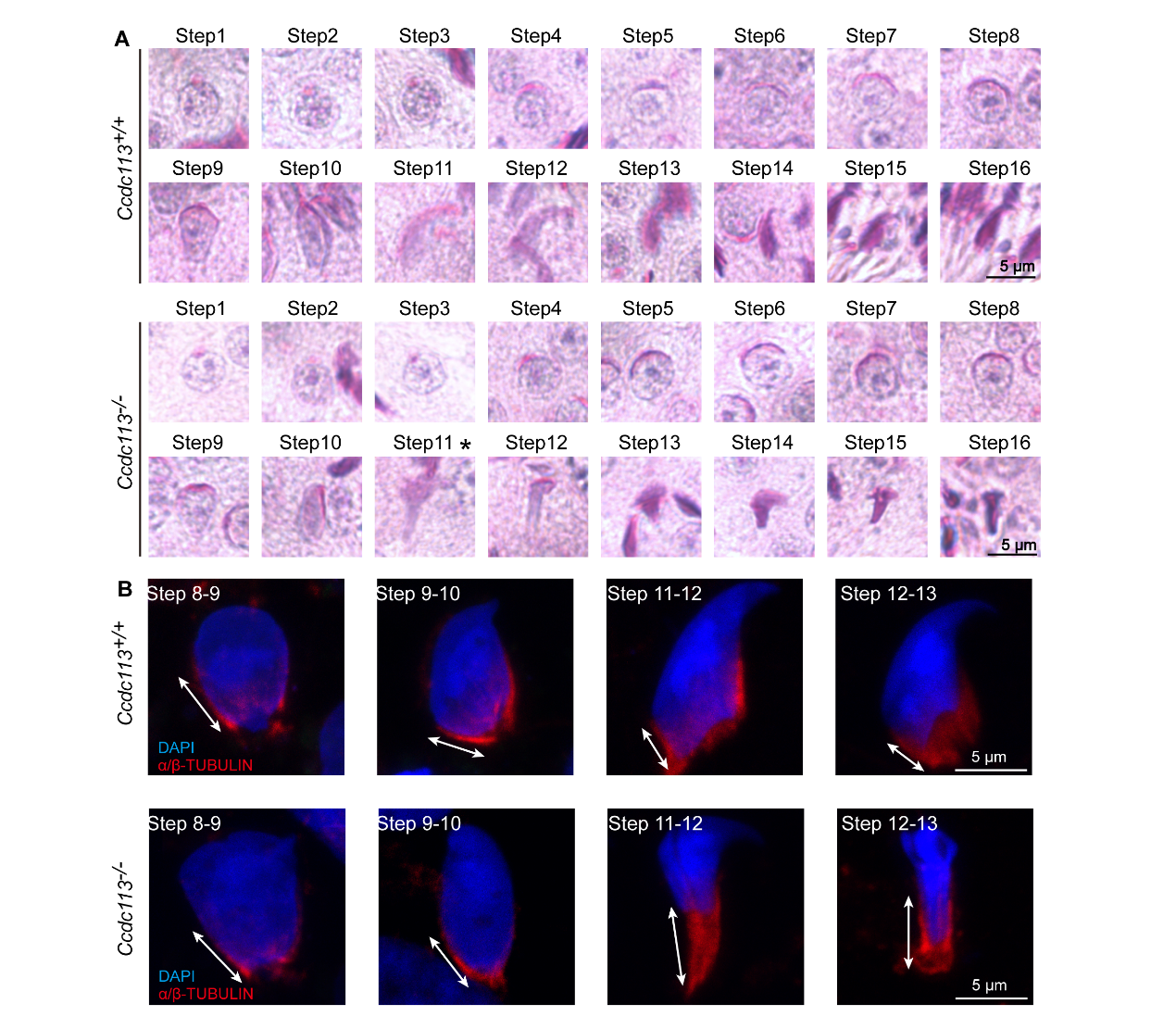

**Figure 3-figure supplement 1. *Ccdc113* knockout leads to abnormal sperm head shaping defect.**  (A) PAS staining of spermatid at different steps from *Ccdc113^+/+^* and *Ccdc113^-/-^* mice. Asterisk indicate abnormal spermatid shapes were found from step 11. (B) Spermatids from different manchette-containing steps were stained with α/β-tubulin antibody (red) to visualize the manchette. Step11 of *Ccdc113^-/-^* spermatids displayed abnormal elongation of the manchette.

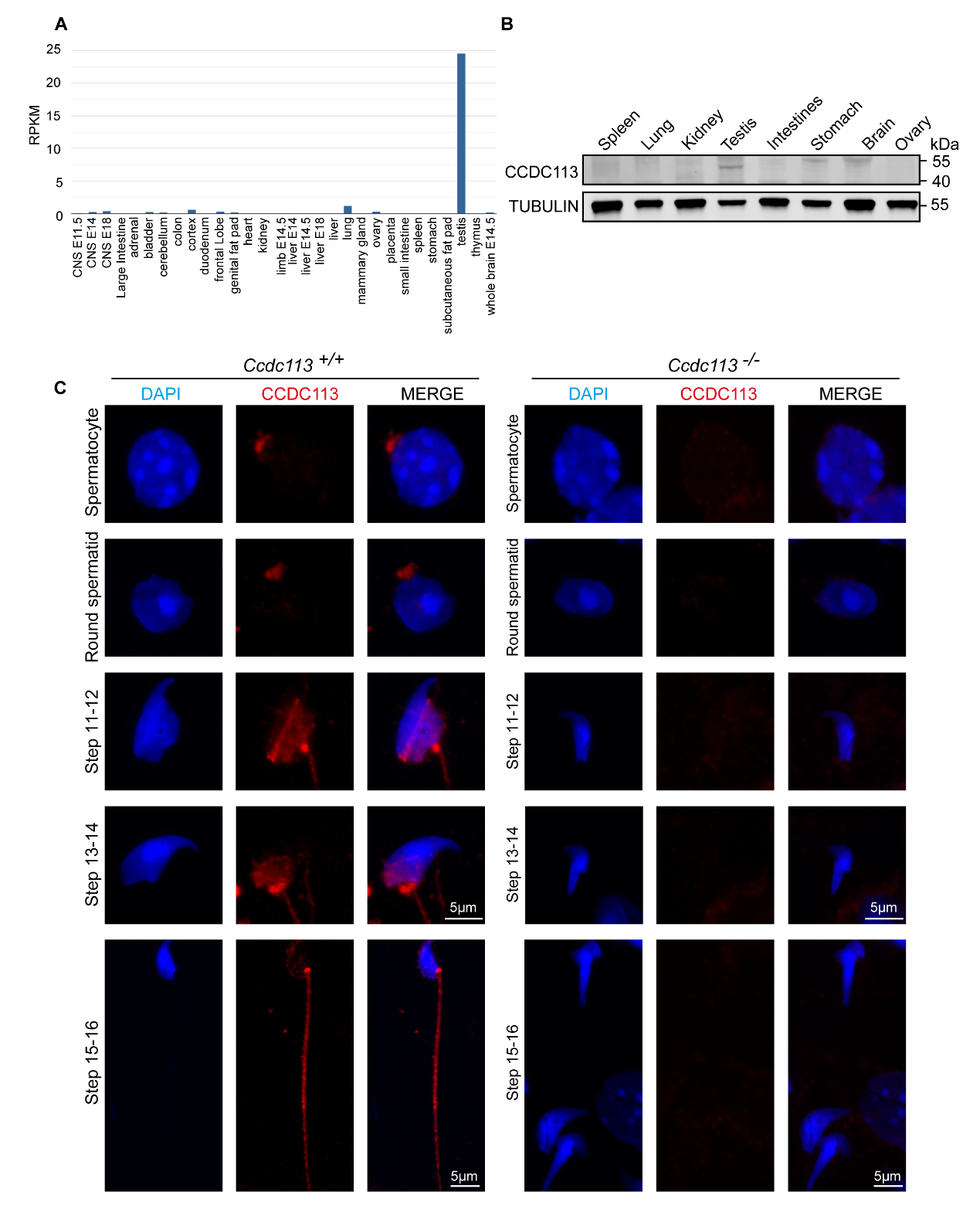

**Figure 4-figure supplement 1. Subcellular localization of CCDC113 in testicular germ cells from *Ccdc113^+/+^* and *Ccdc113^–/–^* mice.** (A) RNA expression data from various tissues using RNA-Seq data from Mouse ENCODE (<https://www.ncbi.nlm.nih.gov/gene/244608>). CCDC113 is highly expressed in the testis, but not significantly in the ovary and brain. (B) CCDC113 was predominately expressed in testis. Immunoblotting of CCDC113 was performed in the spleen, lung, kidney, testis, intestine, stomach, brain and ovary. TUBULIN served as the loading control. (C)The immunofluorescence of CCDC113 in *Ccdc113^+/+^* and *Ccdc113^–/–^* mice. Testicular germ cells were stained with anti-CCDC113 antibody (red), and the nucleus was stained with DAPI.
